# Astrocyte modulation of synaptic plasticity mediated by activity-dependent Sonic hedgehog signaling

**DOI:** 10.1101/2024.04.05.588352

**Authors:** Anh Duc Le, Marissa Fu, Riya Kumar, Gloria Zacharias, A. Denise R. Garcia

## Abstract

The influence of neural activity on astrocytes and their reciprocal interactions with neurons has emerged as an important modulator of synapse function. Astrocytes exhibit activity-dependent changes in gene expression, yet the molecular mechanisms by which they accomplish this have remained largely unknown. The molecular signaling pathway, Sonic hedgehog (Shh), mediates neuron-astrocyte communication and regulates the organization of cortical synapses. Here, we demonstrate that neural activity stimulates Shh signaling in cortical astrocytes and upregulates expression of Hevin and SPARC, astrocyte derived molecules that modify synapses. Whisker stimulation and chemogenetic activation both increase Shh activity in deep layers of the somatosensory cortex, where neuron-astrocyte Shh signaling is predominantly found. Experience-dependent Hevin and SPARC require intact Shh signaling and selective loss of pathway activity in astrocytes occludes experience-dependent structural plasticity. Taken together, these data identify Shh signaling as an activity-dependent, neuronal derived cue that stimulates astrocyte interactions with synapses and promotes synaptic plasticity.

## INTRODUCTION

Activity-dependent gene expression is a fundamental property of neurons by which they modify synapses in response to experience. The mechanisms by which neurons convert activity into gene expression are well characterized and involve Ca^2+^ elevations that trigger downstream molecular signaling programs, including activation of immediate genes such as *Fos* and *Jun*^1^. Emerging evidence demonstrates that astrocytes also exhibit activity-dependent gene expression that promotes synaptic plasticity^2–4^. How astrocytes couple neuronal activity into changes in gene expression is unknown.

Astrocytes are now recognized as influential partners in synaptic function. They associate intimately with synapses, comprising the third partner in the tripartite synapse, along with the pre and postsynaptic terminals^5^. Astrocytes monitor and respond to neural activity^6–9^ and engage in reciprocal regulation of synapse function^10–14^ through a large complement of receptors, transporters, and ion channels embedded within their membranes^15–17^. They also exert broad influence on the plasticity of synapses and circuits in development and in learning and memory^18–22^. Astrocytes deploy various mechanisms to modulate synaptic structure and function, including regulating receptor composition at the synapse^23,24^, release of synaptogenic proteins^25–27^, synaptic pruning^28^, and release of gliotransmitter^29^. Transcriptional profiling shows that astrocytes also undergo activity-dependent changes in gene expression^2–4^. These activity-dependent transcriptional responses are as robust as many classes of neurons^4^ and include several genes associated with synapse formation and function^2,30,31^. Yet, the precise molecular mechanism by which astrocytes convert neuronal activity into gene expression remains unknown.

The molecular signaling pathway, Sonic hedgehog (Shh) is emerging as an important mediator of reciprocal interactions between neurons and astrocytes. In the postnatal and adult brain, SHH is produced by neurons which acts directly on neighboring astrocytes, regulating SHH-dependent gene expression programs^32–34^. Notably, genetic ablation of pathway activity, selectively in astrocytes, decreases expression of the glial-specific, inward rectifying K^+^ channel, K_ir_4.1, perturbing extracellular K^+^ homeostasis and increasing neuronal excitability^35^. Conversely, unrestrained Shh signaling promotes synapse formation mediated by increased expression of synaptogenic cues^34^. These observations point to SHH as a key mediator of neuron-astrocyte interactions that reciprocally shape synapse number and function. Further, non-canonical Shh signaling in neurons is required for establishing cortico-cortical synapses^36^, suggesting diverse and cell-type specific actions of SHH on synaptic regulation. Here, we focus on SHH signaling in astrocytes mediated by canonical transduction of the pathway.

Transduction of Shh signaling is mediated by transcriptional activation of its effectors, the GLI family of transcription factors, that regulate expression of SHH target genes^37^. Binding of SHH to its receptor, Patched (*Ptch1*), relieves inhibition of the obligatory co-receptor, Smoothened (*Smo*), promoting transcription of SHH target genes, including *Gli1*. High levels of SHH promote transcriptional activation of *Gli1*, which acts as a reliable readout of pathway activation^38^. At baseline, transcriptional abundance of *Gli1* in the cortex is low^34^ and its expression is localized to astrocytes in layers 4 and 5, consistent with the distribution of SHH-expressing neurons in layer V^32,36^. However, several components of the pathway, including *Ptch1*, *Gli2,* and *Gli3*, show more widespread expression^32^. Whereas cells expressing *Gli1* at baseline comprise 40-50% of astrocytes in layers 4 and 5^35^, expression of *Ptch1*, *Gli2* and *Gli3* is found in nearly all astrocytes, indicating that many more cells possess the machinery for transduction of SHH signal^32^. This suggests dynamic activity of the pathway in individual cells. What regulates activation of the pathway is not known. High frequency stimulation of cultured hippocampal neurons promotes SHH release that is sensitive to TTX treatment^39^, suggesting that neural activity promotes Shh signaling. Interestingly, SHH protein is localized in axons^40–43^, where it is associated with synaptic vesicles^44^ making it well positioned to couple with neuronal activity. Whether physiological neural activity promotes SHH-mediated interactions between neurons and astrocytes *in vivo,* and how these interactions influence synapses, is not known.

To examine this, we exposed juvenile mice to an enriched somatosensory environment to stimulate active whisking. We show that both sensory and chemogenetic stimulation increase Shh signaling in cortical astrocytes. Exposure to an enriched environment produced an increase in synapse number and a concomitant increase in expression of Hevin and SPARC, matricellular proteins produced by astrocytes that modify synapses. Structural plasticity, and experience-dependent upregulation of Hevin and SPARC, are occluded in astrocyte-specific conditional knock out (CKO) mice in which Shh signaling is selectively abolished, demonstrating a requirement for Shh signaling in astrocyte modulation of synapses. Taken together, these findings demonstrate novel, activity-dependent regulation of Shh signaling and establish SHH as a molecular mechanism by which astrocytes couple neuronal activity to gene expression and modulate synaptic plasticity.

## RESULTS

### Neuronal activity stimulates Sonic hedgehog signaling in vivo

To examine whether neural activity stimulates Shh signaling *in vivo*, we used a chemogenetic approach to selectively excite cortical pyramidal neurons with the excitatory hM3DGq designer receptor exclusively activated by designer drugs (DREADD) (**Figure 1A-B**). We injected AAV8-CamKIIα-hM3DGq-mCherry virus^45^ into the somatosensory cortex of *Gli1^nlacZ/+^* mice which express nuclear lacZ at the *Gli1* locus^38^. Because *Gli1* is a canonical target of Shh signaling, and because SHH is required for *Gli1* expression, transcriptional activation of *Gli1* acts as a reliable readout of Shh signaling^38,46^. After 14 days, we treated mice with medicated drinking water containing the designer drug Clozapine N-Oxide (CNO) for seven days to chronically activate the receptor^47^. Transfected cells expressed *c-Fos* **(Figure 1C)**, and CNO-treated mice showed a significant increase in the number of c-Fos-labeled cells compared to virus-injected animals that were fed unmedicated drinking water, indicating successful activation of neurons (**Figure 1D-E**). We observed a significant increase in the number of βGal+ cells in the somatosensory cortex in the CNO-treated group compared to untreated controls (**Figure 1F-G**). These cells colocalized with Sox9, a well-established marker for identifying astrocytes^48^ (**Figure 1H**). Taken together, these data demonstrate that neuronal excitation stimulates Shh signaling in astrocytes *in vivo*.

**Figure 1.**
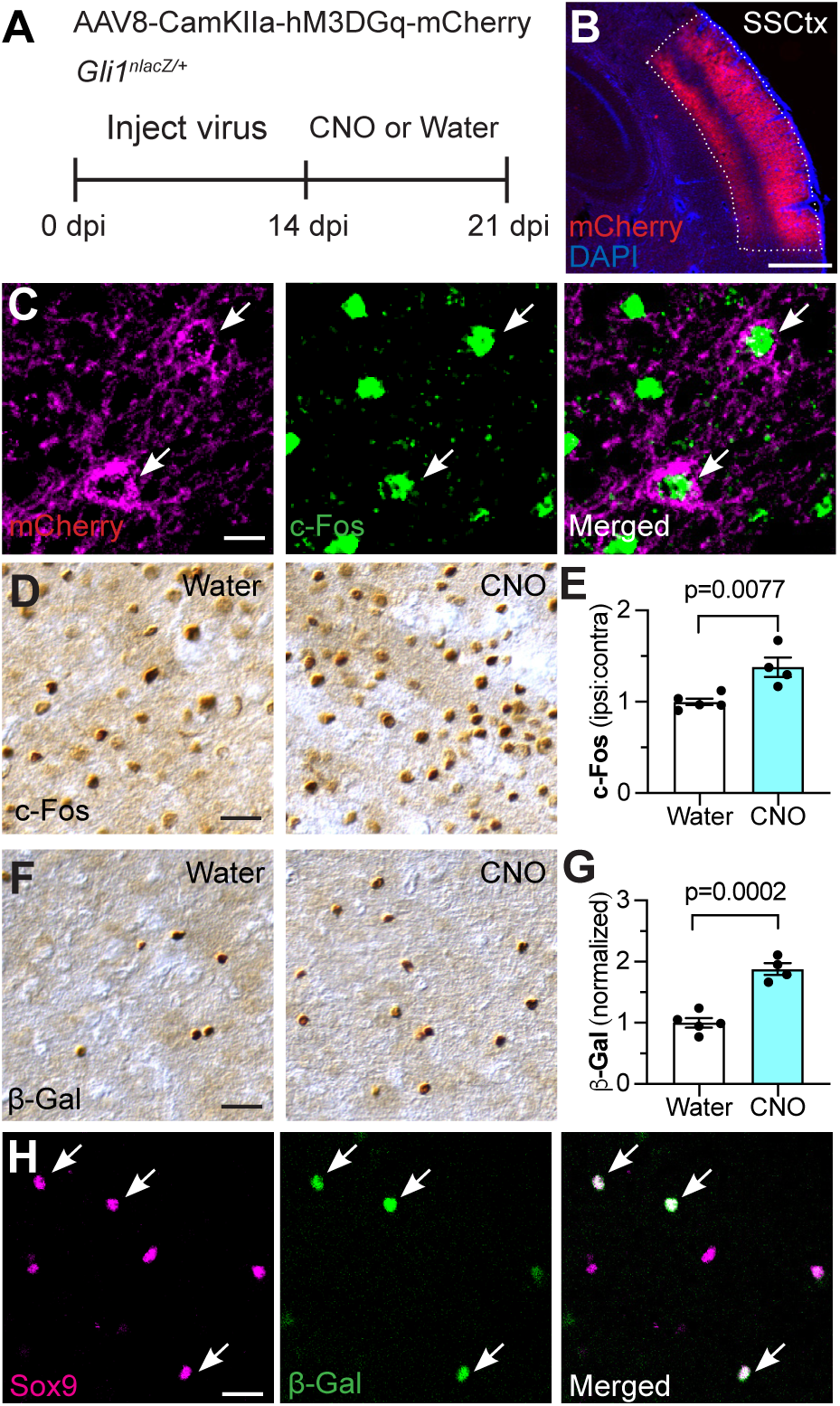
Neuronal activity stimulates Shh signaling. (**A**) Schematic depicting experimental approach. Gli1^nlacZ/+^ mice were injected with AAV8-CamKIIα-hM3DGq-mCherry in the somatosensory cortex. Two weeks later, mice were fed CNO-medicated or unmedicated drinking water for 7 days before analysis. (**B**) Transfected region can be visualized with mCherry. SSCtx, somatosensory cortex. Scale bar, 250 µm. (**C**) mCherry (magenta, left panel) and c-Fos (green, middle panel) in transfected region. Arrows point to cells co-expressing mCherry and c-Fos. Scale bar, 25 µm. (**D, F**) Brightfield immunohistochemistry for c-Fos (D) and βGal (F) in the somatosensory cortex. Scale bar, 25 µm. (**E, G**) Stereological quantification of c-Fos (E) and βGal (G) in the transfected region shows increased neural activity and increased Shh signaling in mice that received CNO compared to water-fed controls. (**H**) Immunofluorescent staining for Sox9 (magenta, left panel) and βGal (green, middle panel) in transfected region. Arrows point to βGal cells co-labeled with Sox9. Scale bar, 25 µm. Data points represent individual animals; bars show mean ± SEM; Student’s t-tests.

### Sensory experience stimulates Sonic hedgehog signaling

We next examined whether a more physiological stimulus can increase Shh signaling. To test this, we placed *Gli1^nlacZ/+^* mice in an enriched sensory environment designed to stimulate active whisking^49^. Mice were weaned into large rat cages from which strings of beads were hung from the wire cage lid in a manner requiring constant interaction at postnatal day 21 (P21). Control mice were weaned into similar rat cages with only standard nesting materials and no beads. Mice were housed in the enriched environment (EE) or standard housing (SH) conditions for three weeks (**Figure 2A**). A significant increase in the number of cells labeled with cFos was observed in the somatosensory cortex of mice housed in EE compared to SH conditions (**Figure 2C**), confirming increased activity in this region. Mice housed in EE showed a significant increase in the number of βGal-labeled cells compared to control mice housed in SH conditions (**Figure 2B, D**), indicating an increase in Shh activity in response to sensory experience. Notably, there was no difference in either cFos or βGal labeling between EE and SH conditions in the visual cortex (**Figure 2E-F**), demonstrating that sensory stimulation does not produce a generalized increase in Shh signaling but rather, selectively increases activity of the pathway in circuits that are stimulated by experience.

**Figure 2.**
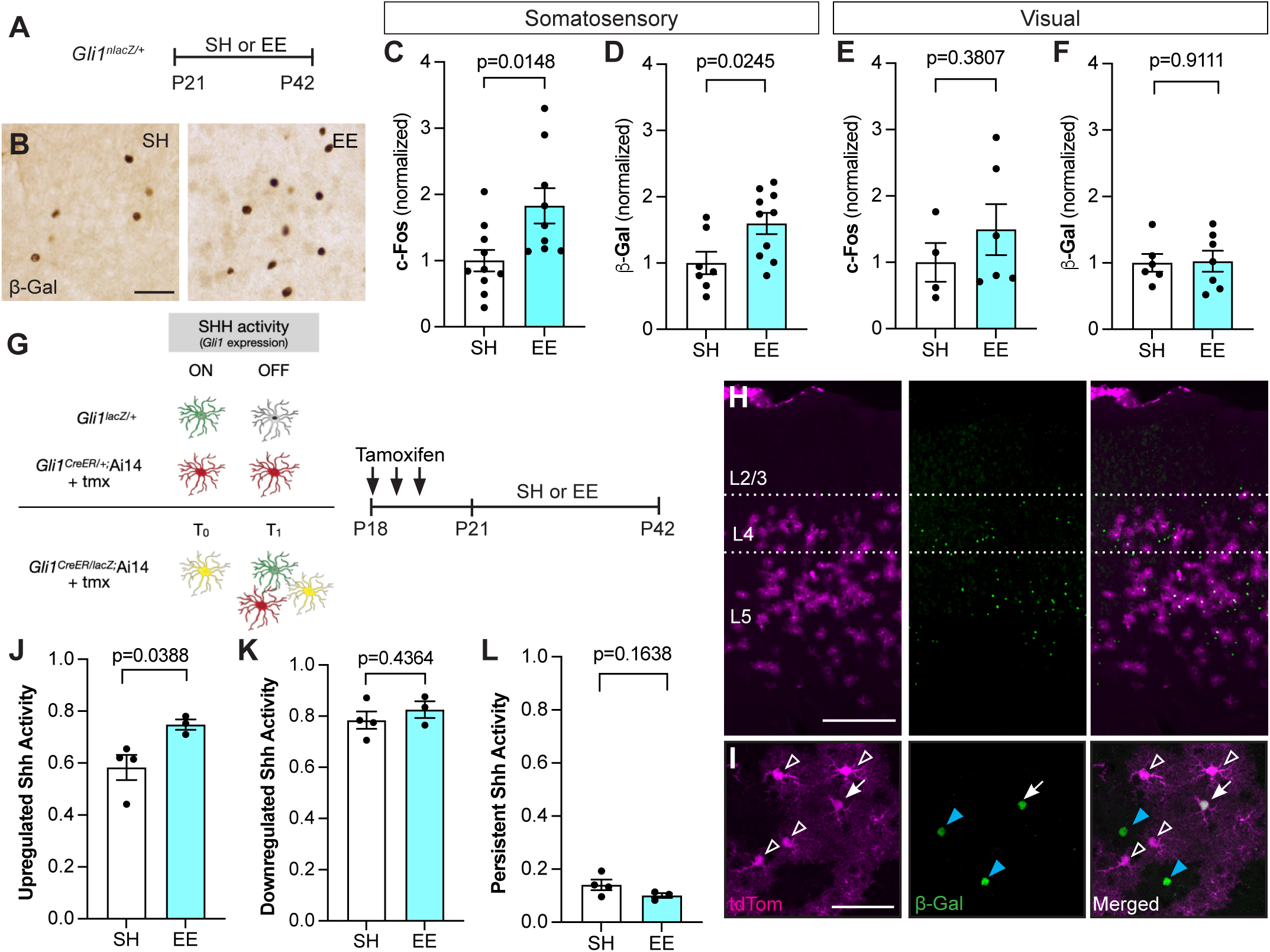
Enriched experience stimulates Shh signaling. **(A)**Experimental timeline. *Gli1^nlacZ/+^*mice were weaned into standard housing (SH) or enriched experience (EE) cages at P21 and analyzed at P42. **(B)** Representative brightfield immunohistochemistry for βGal in the somatosensory cortex. Scale bar, 50 μm. **(C-F)** Stereological quantification of cells expressing the immediate-early gene *c-Fos* (C, E) or βGal (D, F) in the somatosensory (C, D) or visual (E, F) cortices. **(G)** Schematic depicting genetic strategy to label cells with temporally distinct Shh signaling (left panel) and timeline of experiment (right panel). In *Gli1^nlacZ/+^*mice, βGal expression labels cells with active Shh signaling. In *Gli1^CreER^;*Ai14 mice, Cre-mediated recombination promotes permanent expression of tdTom. In *Gli1^CreER/nlacZ^;* Ai14 mice, tdTom expression identifies cells with Shh signaling at baseline (t_0_) and βGal expression identifies Shh signaling in cells at the end of the experiment (t_1_). Double labeled cells identify cells with persistent Shh signaling throughout the experiment. **(H-I)** Low power (H) and high power (I) images showing tdTom expression (magenta, left panel) and immunostaining for β-Gal (green, middle panel; merged image, right panel) in the cortex of mice housed in EE as described in (G). Scale bar, 250 μm. Individual cells show those with upregulation (blue arrowhead), downregulation (open arrowhead) and persistent (white arrow) Shh activity. Scale bar, 50 μm. **(J-L)** The fraction of βGal-labeled cells that are single labeled (J), tdTom-labeled cells that are single labeled (K), and all cells that are double labeled (L) in SH or EE. Data points represent individual animals; bars show mean ± SEM; Student’s t-tests.

Colocalization with cell-type specific markers including Sox9, CAII, and NeuN to identify astrocytes, oligodendrocytes, and neurons, respectively, showed that 96% of βGal-labeled cells correspond to astrocytes in both SH and EE (**Supplemental Figure 1**) consistent with our previous observation that astrocytes are the primary targets of canonical SHH signaling in the adult brain^32^. βGal+ cells did not express Ki-67 (**Supplemental Figure 1**), ruling out the possibility that the observed increase in cell number is due to proliferation. Because sensory activity stimulates Shh signaling, we next asked whether sensory deprivation lowers activity of the pathway. To do this, we performed unilateral whisker trimming on P21 *Gli1^nlacZ/+^* mice for three weeks, starting at birth. We analyzed the deprived, contralateral hemisphere and compared it to the intact, ipsilateral hemisphere. We observed significantly fewer βGal+ cells in the deprived, compared to the intact, hemisphere (**Supplemental Figure 2**). Analysis of the auditory cortex showed whisker trimming had no effect on Shh signaling in this region (**Supplemental Figure 2**). Taken together, these data demonstrate that activation of distinct sensory circuits selectively stimulates Shh signaling in cortical astrocytes in a circuit-specific manner.

To precisely identify cells that show increased Shh activity following experience, we combined the direct reporter approach with a tamoxifen-dependent labeling strategy, enabling us to monitor temporally distinct populations of cells experiencing Shh signaling. We crossed *Gli1^nlacZ/+^* mice with *Gli1^CreER/+^*^;^Ai14 mice carrying the Cre-dependent Ai14 reporter allele expressing tdTomato (tdTom) (*Gli1^nlacZ/CreER^*;Ai14), enabling identification of cells with Shh signaling at the start and conclusion of sensory enrichment. Because *lacZ* expression is directly regulated by transcriptional activity of *Gli1*, cells actively transducing SHH can be monitored by immunohistochemistry for βGal, the protein product of the *lacZ* gene. In contrast, tamoxifen-dependent, Cre-mediated recombination produces permanent expression of tdTom in cells, even after Shh activity is downregulated (**Figure 2G**). Thus, in *Gli1^nlacZ/CreER^*;Ai14 mice, *lacZ* expression reflects active Shh signaling whereas tamoxifen-dependent Cre-mediated expression of tdTom reflects historical signaling. Although these mice are effectively *Gli1* nulls, GLI1 is dispensable for Shh signaling due to redundant activator function of GLI2^38,50^ and we did not observe any gross anatomical or behavioral phenotypes, consistent with previous studies^38,51^. To mark cells with Shh signaling at baseline, mice received tamoxifen before placing them in EE or SH housing for three weeks (**Figure 2G**). We first examined the population of βGal+ cells and found a large fraction of cells that were negative for tdTom (**Figure 2H, I**), indicating Shh activity in a population of cells distinct from those marked at the start of the experiment. Notably, there was a significant increase in single-labeled βGal cells after EE compared to SH (**Figure 2J**). Whereas 59% of βGal labeled cells were single labeled in SH mice, this fraction increased to 76% in EE mice (**Figure 2J**) demonstrating experience-dependent activation of the pathway in cells distinct from those at baseline. These cells were found in layers 4 and 5 (**Figure 2H**), suggesting that experience-dependent Shh activity remains localized to deep cortical layers, consistent with the localization of SHH-expressing neurons in layer V^32,36^. We next examined the population of cells expressing tdTom. tdTom+ were localized primarily in layers 4 and 5 and show a bushy morphology characteristic of protoplasmic astrocytes, consistent with previous studies^32,35^ (**Figure 2H-I**). In SH mice, a large proportion of tdTom+ cells did not colabel with βGal (**Figure 2I, K**), indicating downregulation of pathway activity. This fraction did not change after exposure to EE (**Figure 2K**). We also observed a third population of cells, those co-expressing both tdTom and βGal, suggesting either persistent or recurrent Shh activity in these cells (**Figure 2I, L**). This fraction was similar between SH and EE conditions (**Figure 2L**). These data suggest that SHH activity in individual cells is highly dynamic and that over a three-week period, individual cells are not likely to experience persistent SHH signaling.

### Sensory experience dynamically modulates Sonic hedgehog signaling

We next examined whether the experience-induced increase in Shh activity reflects a long-term change in Shh signaling. We first examined whether a much shorter time frame was sufficient to increase Shh activity. We housed *Gli1^nlacZ/+^* mice in either SH or EE for two days starting at P21. Mice housed in EE showed a significant increase in βGal+ cells compared to SH controls (**Figure 3A-B**), suggesting that brief exposure to sensory enrichment is sufficient to stimulate Shh signaling. To examine whether experience-dependent Shh signaling persists after removal of the stimulus, we housed *Gli1^nlacZ/+^* mice in EE for 2 days, then housed them in SH conditions for 2 days and compared *Gli1* activity with mice continuously housed in EE or SH conditions for 4 days. To rule out the possibility that any observed differences in βGal number may reflect perdurance of the protein rather than sustained Shh activity, we used single molecule fluorescent *in situ* hybridization (smFISH) by RNAScope to more directly measure *Gli1* transcripts (**Figure 3C**). In mice continuously housed in EE, we found an increase in cells expressing *Gli1* compared to SH control mice (**Figure 3C-D**). However, in mice returned to SH after 2 days of EE, *Gli1* expression returned to baseline (**Figure 3C-D**). Notably, individual cells expressing *Gli1* also showed a dramatic increase in the number of transcripts in the nucleus after exposure to EE compared to SH-housed mice (**Figure 3C**). This suggests that neural activity not only stimulates Shh signaling in new cells, but also further increases activity of the pathway in cells with baseline Shh signaling. These data demonstrate that Shh signaling is highly dynamic and responsive to changing experiences.

**Figure 3.**
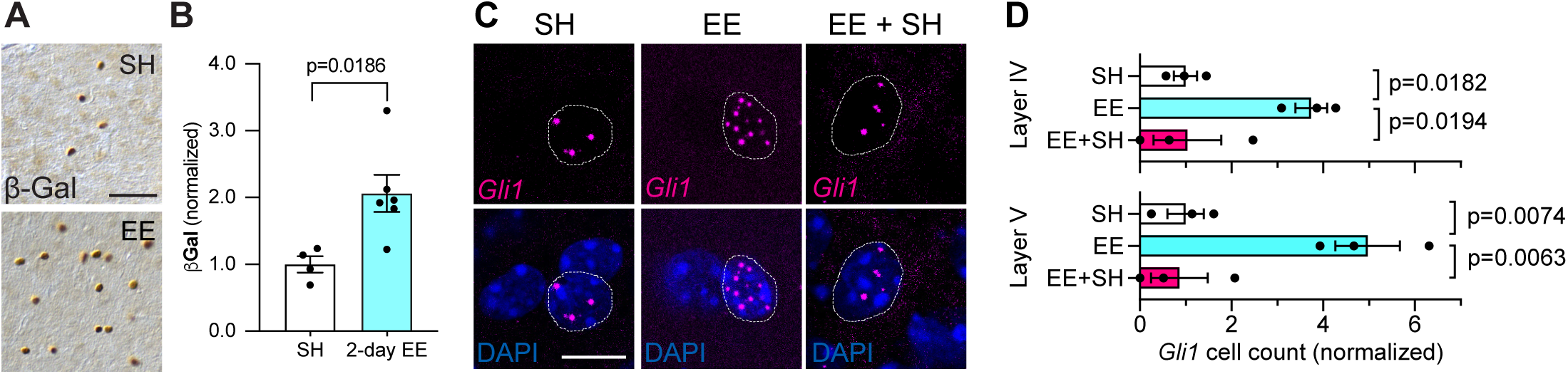
Shh signaling responds rapidly to sensory activation and is not persistent. (**A**) Brightfield immunohistochemistry for β-Gal in the somatosensory cortex of *Gli1^nlacZ/+^* mice housed in SH (upper panel) or EE (lower panel) for 2 days. Scale bar, 50 μm. (**B**) Stereological quantification of β-Gal shows increased SHH activity in the somatosensory cortex after 2 days of EE. (**C**) Representative images of fluorescent *in situ* hybridization for *Gli1* mRNA in deep layers of the cortex from SH, EE and EE+SH housing conditions. Dotted line outlines individual DAPI nuclei with *Gli1* mRNA. Note the increase in *Gli1* transcripts in EE compared to SH. Scale bar, 10 µm. (**D**) *Gli1*+ cell count in Layers IV and V of the cortex from mice housed in SH or EE for 4 days or EE for 2 days followed by SH for 2 days (EE+SH). In each layer, 100 - 800 DAPI cells analyzed per animal. Data points represent individual animals; data represent mean ± SEM; Student’s t-test (B) and one-way ANOVA with Tukey’s multiple comparisons (D).

### Sensory experience does not increase the number of Shh-expressing neurons

In the cortex*, Shh* is expressed predominantly in a subset of excitatory pyramidal neurons in layer V^35,36^. To determine whether sensory experience increases the number or distribution of *Shh*-expressing cells, we housed *Shh^CreER/+^*; Ai14 mice in EE or SH conditions. Mice received tamoxifen over the last three days of the housing experience and were analyzed two weeks later (**Figure 4A**). Labeled neurons were found primarily in layer V and we found no difference in the number of *Shh*-expressing neurons between SH and EE (**Figure 4B-E**). Because astrocytes in layer IV show high levels of Shh activity, we also examined the ventral posterior medial nucleus of the thalamus (VPM) where thalamocortical neurons projecting to layer IV of the barrel cortex reside^52^. We detected few, if any, labeled neurons in this region (**Figure 4D**) suggesting that Shh signaling in cortical astrocytes is initiated by ligand released from local neurons. We also performed chemogenetic activation of cortical neurons in *Shh^CreER/+^*; Ai14 mice. We injected mice with AAV5-Syn-hM3DGq-HA virus and began CNO treatment two weeks later. Animals received CNO or unmedicated water for 10 days. To mark *Shh*-expressing cells, animals received tamoxifen over the last three days of CNO treatment and were analyzed two weeks later (**Figure 4F**). In agreement with our environmental stimulation, there was no difference in the number or distribution of *Shh*-expressing cells between the CNO treated and control animals (**Figure 4G-J**). These data suggest that the increase in astrocytic signaling reflects an increase in availability of the ligand, rather than an increase in the number of neurons expressing *Shh*. This may arise from an increase in SHH release or from increased diffusion in the extracellular space. This suggests that while transduction of Shh signaling in astrocytes is dynamically modulated between individual cells, SHH ligand is derived from a defined population of neurons.

**Figure 4.**
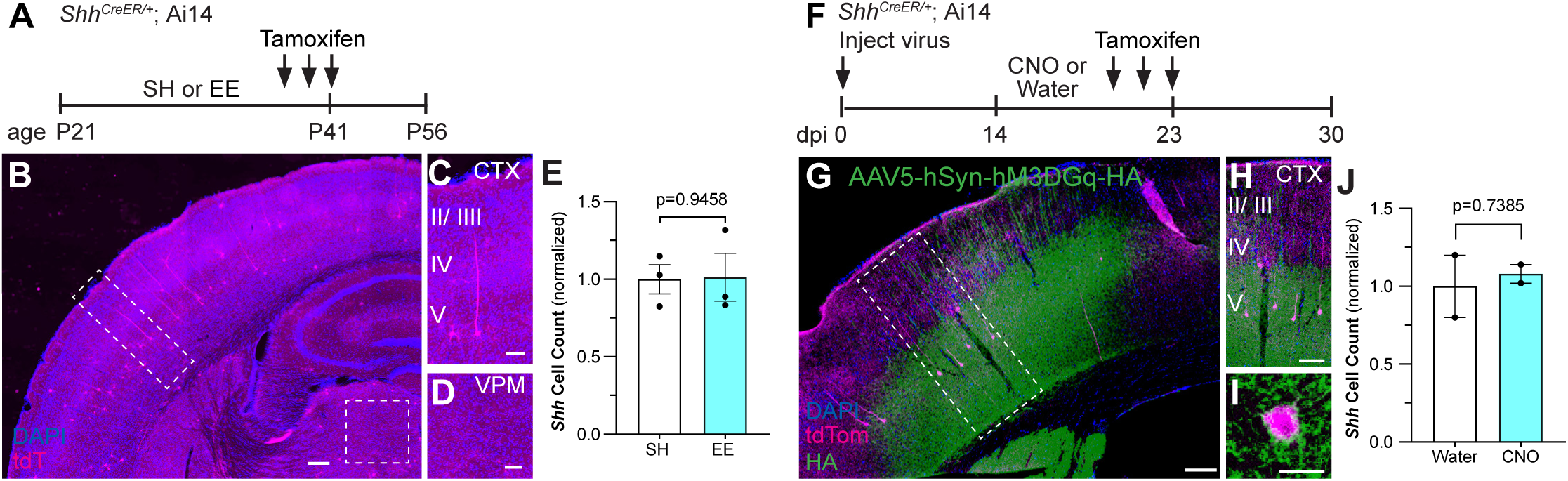
Sensory experience does not increase the number of *Shh*-expressing neurons. (**A**) Schematic depicting experimental approach. P21 *Shh^CreER/+^;* Ai14 mice were weaned into SH or EE cages and received 3 doses of tamoxifen from P39 - P41. Tissues were analyzed two weeks after the last tamoxifen dose. (**B-D**) Low power image showing tdTom labeled cells (magenta) in *Shh^CreER/+^;* Ai14 mice. Labeled cells are found predominantly in layer V. Insets show individual cells with neuronal morphology in the cortex (C) and no labeled cells in the ventral posterior medial (VPM) nucleus of the thalamus (D). Counterstained with DAPI (blue). Scale bar, 25 μm. (**E**) The number of *Shh* neurons in the cortex in SH and EE mice. (**F**) *Shh^CreER/+^*; Ai14 mice were injected with AAV5-hSyn-hM3DGq-HA targeting the somatosensory cortex. After 2 weeks, mice were fed CNO-medicated or unmedicated water for 10 days, receiving 3 doses of tamoxifen during the last 3 days. Tissues were analyzed seven days later. dpi, days post injection. (**G-I**) Immunolabeling for HA (green) shows the transduced region overlapping with tdTom labeled (magenta) cells in the cortex. Inset shown in (H). Individual cell shown at high power in (I). Scale bar, 25 μm. (**J**) The number of *Shh* neurons in the cortex between water control versus CNO. Data points represent individual animals; bars show mean ± SEM; Student’s t-tests.

### Astrocytic Sonic hedgehog signaling is required for deep layer synaptic plasticity

Sensory experience promotes structural plasticity of cortical synapses^49^. To determine whether experience-dependent Shh signaling in astrocytes promotes synaptic plasticity, we examined dendritic spines of layer V cortical neurons. Excitatory synapses are localized predominantly on dendritic spines^53^, which demonstrate remarkable plasticity in response to sensory experience^49,54–57^. To determine whether Shh signaling plays a role in astrocyte modulation of experience-dependent plasticity, we examined spine density of cortical neurons in conditional knock out (CKO) mice lacking *Smo,* the obligatory co-receptor for transduction of Shh signaling, selectively in astrocytes. We used *Gfap-Cre*; *Smo^fl/fl^* mice (*Gfap Smo* CKO) in which *Smo* is deleted in *Gfap*-expressing cells, effectively abolishing Shh signaling in all astrocytes beginning at birth^32,35^. These mice also carry a *Thy1*-GFPm allele which labels a subset of layer V pyramidal neurons^58^, enabling the visualization of individual dendrites and spines. *Gli1* expression measured by qPCR showed a 90% reduction in *Gfap Smo* CKO mice compared to WT littermate controls lacking Cre, confirming effective ablation of Shh activity (**Supplemental Figure 3**). We analyzed apical dendrites of layer V pyramidal neurons in the barrel cortex, focusing on dendritic segments in layers 4 and 5 where high levels of Shh activity are found. WT (Cre-negative littermates) mice housed in EE conditions showed a significant increase in the density of protrusions compared to control SH mice (**Figure 5A,C**), demonstrating that enriched sensory experience promotes structural synaptic plasticity, consistent with previous studies^49,56^. We previously demonstrated that *Gfap Smo* CKO mice showed an elevated spine density arising from deficits in developmental elimination of synapses^35^. Here, we find that sensory enrichment did not produce any further increases in spine density (**Figure 5B,C**), consistent with a requirement for astrocytic Shh signaling in structural plasticity. Alternatively, this may instead reflect a limit beyond which no further spines can be added. Shh signaling acts on astrocyte progenitor cells in the perinatal subventricular zone that generate cortical astrocytes^51^. Because recombination begins at P0 in cortical astrocyte progenitors in this *Gfap-Cre* line^32^, this impaired plasticity may reflect developmental perturbation of cortical astrocyte development.

**Figure 5.**
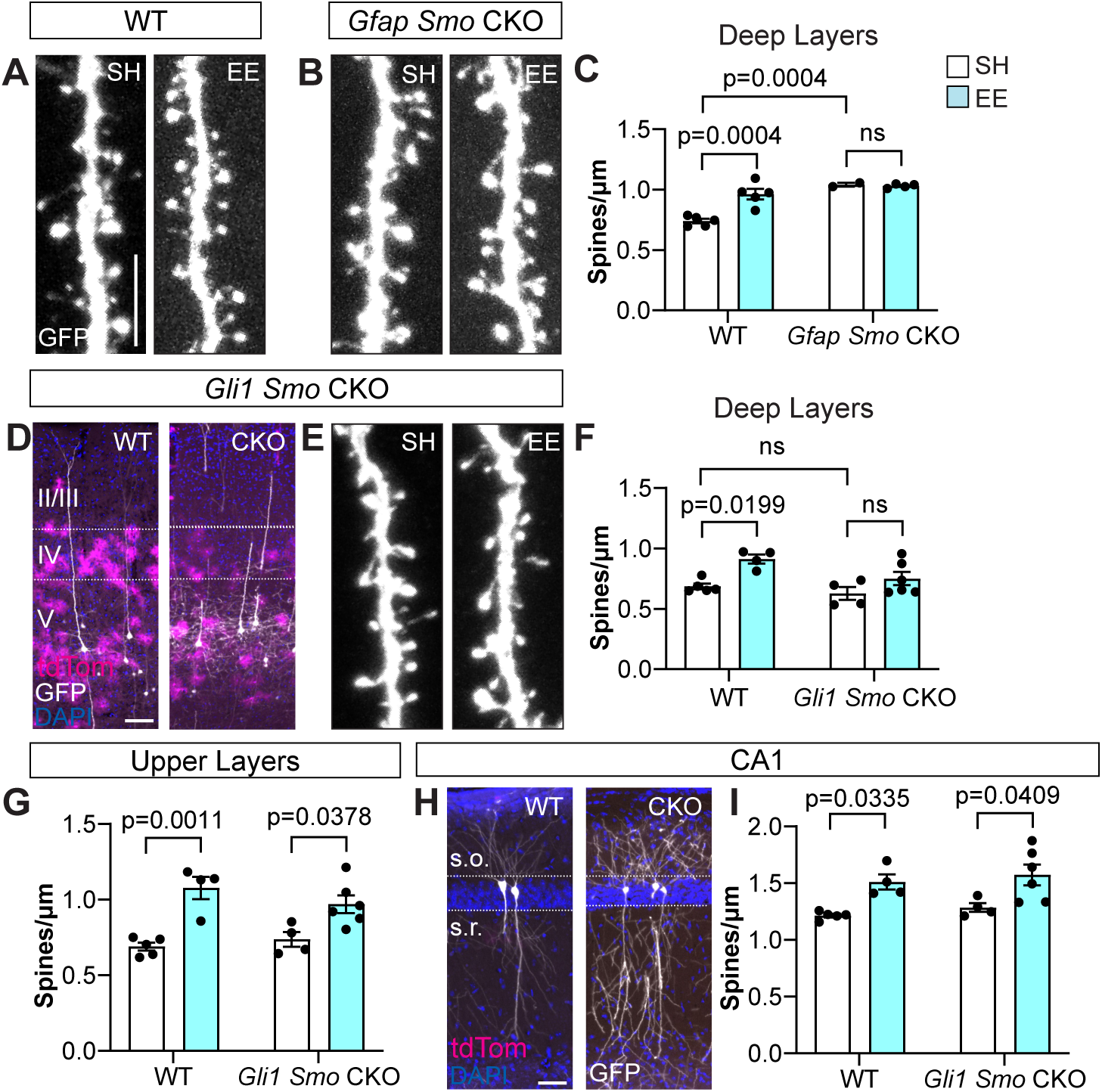
Experience-dependent structural plasticity requires Shh signaling. (**A-B**) Representative dendritic segments in deep layers of WT (A) and *Gfap Smo* CKO (B) mice housed in SH or EE. Scale bar, 5 μm. (**C**) Spine density of dendritic segments in deep layers of WT and *Gfap Smo* CKO mice housed in SH or EE. (**D**) tdTom (magenta) labeled astrocytes and GFP (white) labeled neurons, counterstained with DAPI (blue), in the cortex of WT (left) and *Gli1 Smo* CKO (right) mice. Scale bar, 100 μm. (**E**) Dendritic segments from *Gli1 Smo* CKO mice housed in SH or EE. (**F**) Spine density of deep layer dendritic segments from WT and *Gli1 Smo* CKO mice housed in SH or EE. (**G**) Spine density of upper layer dendritic segments from WT and *Gli1 Smo* CKO mice housed in SH or EE. (**H**) tdTom (magenta) and GFP (white) labeled cells in area CA1 of the hippocampus from WT and *Gli1 Smo* CKO mice, counterstained with DAPI (blue). Area CA1 contains few tdTom labeled cells reflecting the absence of Shh activity in hippocampal astrocytes. Scale bar, 50 μm. (**I**) Spine density from apical dendrites of CA1 neurons in WT and *Gli1 Smo* CKO mice housed in SH or EE. Data points represent individual animals, bars show mean ± SEM, *n ≥ 3 - 5* dendritic segments analyzed per animal. Statistical testing performed by two-way ANOVA with Tukey’s multiple comparisons. s.o., stratum oriens; s.r., stratum radiatum.

To more directly test whether Shh signaling in mature astrocytes is required for synaptic plasticity, we generated *Gli1^CreER/+^*; *Smo^fl/fl^*; Ai14 mice (*Gli1 Smo* CKO), in which tamoxifen-dependent *Smo* deletion enables temporal control. Tamoxifen administration at P14 or older limits *Smo* deletion to postmitotic, *Gli1* expressing cells primarily in layers 4 and 5 (**Figure 5D**)^51^. *Gli1 Smo* CKO mice received tamoxifen over three days between P18 – P20 and were housed in EE or SH at P21 for two days. We measured *Gli1* expression of cortical tissues by qPCR and detected a 66% reduction in *Gli1* transcripts in *Gli1 Smo* CKO mice compared to WT littermate controls that were Cre-negative, confirming effective interruption of Shh signaling (**Supplemental Figure 3**). Cre-negative littermates with and without tamoxifen as well as Cre-positive littermates without tamoxifen showed no significant difference in dendritic spine density and were subsequently pooled as WT controls (**Supplemental Figure S3**). WT mice housed in EE showed a significant increase in spine density compared to SH controls (**Figure 5F**). In contrast, *Gli1 Smo CKO* mice failed to show a significant difference in spine density between housing conditions, demonstrating a requirement for astrocytic Shh signaling in structural plasticity (**Figure 5E,F**). To determine whether plasticity is perturbed globally, we analyzed regions where Shh activity in astrocytes is low or absent. Notably, analysis of dendritic segments in upper layers (layers II/ III) of both WT and *Gli1 Smo CKO* mice showed a significantly higher spine density in EE compared to SH (**Figure 5G**), suggesting that structural plasticity of these synapses is intact. We also analyzed apical dendrites of pyramidal neurons in area CA1 of the hippocampus, which lack astrocytic Shh activity^35^ (**Figure 5H**). *Gli1 Smo CKO* mice housed in EE also showed a significant increase in spine density compared to SH (**Figure 5I**), further supporting the specificity of this mouse model. Taken together, these data demonstrate that Shh signaling in astrocytes promotes structural plasticity induced by enriched sensory experience. The observation that plasticity is impaired specifically in synapses found where astrocytic Shh activity is high suggests that SHH-dependent astrocyte modulation of synaptic plasticity is mediated by local interactions.

### Sonic hedgehog signaling upregulates expression of astrocyte-derived synapse-modifying cues

In the developing neural tube, Shh acts on neural progenitor cells to regulate the expression of genes that drive morphogenesis and cell type specification. In astrocytes, several lines of evidence demonstrate that Shh regulates genes associated with synapse formation and function, including *Sparc, Kcnj10*, and *Gria1*^33–35^. The observation that structural plasticity is impaired in *Gli1 Smo* CKO mice suggests that Shh regulates an astrocyte-derived synaptogenic cue. Hevin is a matricellular protein belonging to the SPARC family of proteins that is expressed in astrocytes and promotes synapse formation^26,59–61^. To examine whether experience-dependent Shh activity regulates *Hevin*, we examined its expression in the cortex of *Gli1^CreER/+^*;Ai14 mice. Hevin immunostaining in the cortex shows an enrichment in layers IV and V, where Shh activity is highest (**Supplemental Figure 4**). Single cell analysis of deep and upper layer astrocytes in *Gli1^CreER/+^*;Ai14 mice revealed that astrocytes in layers IV and V, identified by tdTom labeling, express higher levels of Hevin compared to those in layer II/ III, identified by Sox9 (**Figure 6A-B**), suggesting that Shh signaling regulates its expression. To test this, we measured *Hevin* from whole cortical lysates of *Gfap Smo* CKO mice by qPCR and found a significant reduction in expression compared to WT controls (**Figure 6C**). We next examined secreted acidic cysteine rich glycoprotein (SPARC), a structurally related protein to Hevin that is secreted by astrocytes and which also acts to modify synapses *in vitro* and *in vivo*^61,62^. Expression of *Sparc* is upregulated in *Glast Ptch1* CKO mice in which Shh signaling is elevated in astrocytes^34^. Similar to Hevin, SPARC expression was enriched in deep layer, relative to upper layer, astrocytes in WT mice (**Figure 6D-E, Supplemental Figure 4**). There was a significant reduction in *Sparc* expression in *Gfap Smo* CKO mice compared to WT controls (**Figure 6F**), demonstrating a requirement for Shh signaling in regulating its expression. Neither *Hevin* nor *Sparc* expression were completely abolished in *Gfap Smo* CKO mice, suggesting that Shh signaling may cooperate with additional mechanisms to regulate their expression. Indeed, although their expression is higher in deep layers of the cortex, their expression is not absent in upper layers (**Figure 6A, D, Supplemental Figure 4**). Taken together, these data show that *Hevin* and *Sparc* are enriched in deep layer astrocytes and further demonstrate a requirement for Shh signaling in regulating their expression.

**Figure 6.**
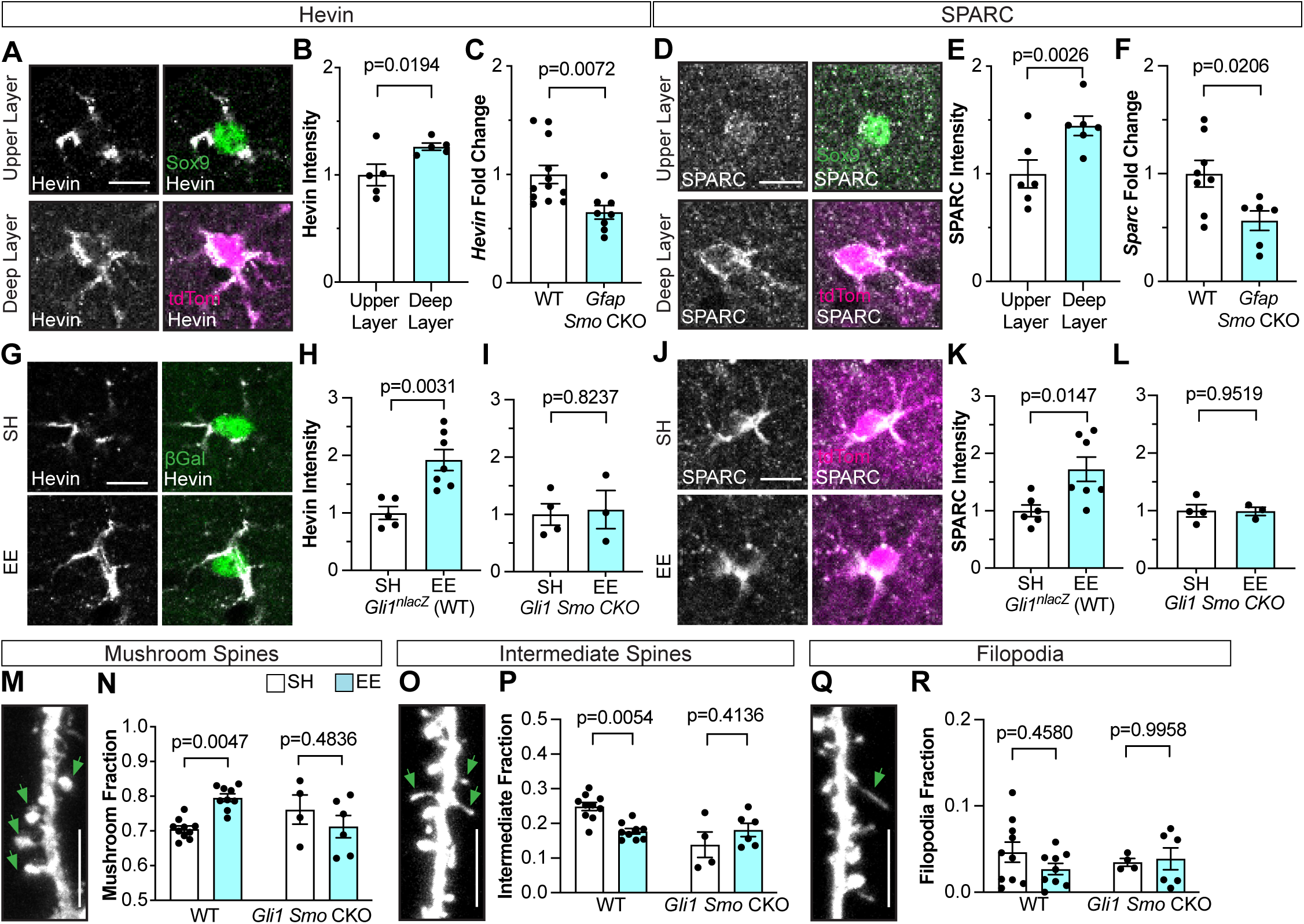
Experience-dependent Hevin and SPARC are Shh-dependent. (**A, D**) Immunofluorescent staining for Hevin (A, gray) and SPARC (D, gray) in individual astrocytes from upper layer labeled with Sox9 (green) and deep layers labeled with tdTom (magenta) in the cortex of P60 *Gli1^CreER/+^;* Ai14 mice. Merged images in right panels. Scale bar, 10 µm. (**B, E**) Analysis of fluorescent intensity of Hevin (B) and SPARC (E) immunostaining in individual astrocytes from upper versus deep layers. *n = 5-6* animals, *≥* 30 cells analyzed per animal. (**C, F**) qPCR for *Hevin* (C) or *Sparc* (F) expression in WT vs *Gfap Smo* CKO mice. *n=8-12* WT mice, *n=6-8 Gfap Smo* CKO mice. (**G**) Immunofluorescent staining for Hevin (gray) and βGal (green) in *Gli1^nlacZ/+^*(WT) mice housed in SH vs EE from P21-P23. Merged images in right panels. Scale bar, 10 μm. (**H, I**) Single cell analysis of Hevin fluorescent intensity in WT (H) and *Gli1 Smo* CKO (I) mice housed in SH vs EE. 25-50 cells analyzed per animal. (**J**) Immunofluorescent staining for SPARC (gray) in tdTom cells (magenta) in *Gli1^CreER/+^; Smo^fl/fl^;* Ai14 (*Gli1 Smo* CKO) mice housed in SH vs EE. Merged images in right panels. Scale bar, 10 μm. (**K, L**) Single cell analysis of SPARC fluorescent intensity in WT (K) and *Gli1 Smo* CKO (L) mice housed in SH vs EE. 25-50 cells analyzed per animal. (**M, O, Q**) Representative examples of dendritic protrusions categorized as mushroom spines (M), intermediate spines (O) and filopodia (Q). Scale bar, 5 μm. (**N, P, R**) Fraction of mushroom spines (N), intermediate spines (P) and filopodia (R) from WT and *Gli1 Smo* CKO mice housed in SH vs EE. At least 300 spines from 3 dendritic segments analyzed per animal. Data points represent individual animals; bars show mean ± SEM; Student’s t-tests (B, C, E, F, H, I, K, L) and 2-way ANOVA with Tukey’s multiple comparisons (N, P, R).

We next asked whether experience-dependent Shh activity promotes an increase in expression of these synapse modifying cues. We performed immunostaining for Hevin and SPARC in *Gli1^nlacZ/+^* mice housed in SH or EE from P21-P23 and analyzed their expression selectively in βGal+ cells in deep layers of the cortex. Mice housed in EE showed a significant increase in Hevin and SPARC compared to SH controls (**Figure 6G, H, K**), demonstrating that enriched sensory experience promotes upregulation of these proteins. Notably, exposure to EE failed to increase Hevin or SPARC in both *Gfap Smo CKO* and *Gli1 Smo CKO* mice (**Figure 6I, L, Supplemental Figure 4**), demonstrating a requirement for Shh signaling in experience-dependent upregulation of these genes.

Hevin and SPARC act, respectively, to modify synapses structurally and functionally. SPARC has been shown to increase the number of GluA1 containing AMPA receptors at synapses^63^, whereas Hevin promotes the formation of spines with a mushroom morphology^61^. Analysis of GluA1 in dendritic spines showed that only a small fraction of spines contained GluA1 receptors and this did not change following exposure to EE (**Supplemental Figure 5**). We found no difference in the fraction of spines containing GluA1 AMPARs in either Gfap Smo CKO or Gli1 Smo CKO mice, arguing against a role for Shh signaling in synaptic GluA1 (**Supplemental Figure 5**).

We next analyzed spine morphology, classifying them into mushroom, thin, and filopodial morphologies and examined whether the proportion of spines in each class change in response to experience and Shh signaling. Spine morphology is associated with synapse size and strength^64,65^. In WT mice housed in EE, we observed an increase in the proportion of spines with a mushroom morphology (**Figure 6M-N**), consistent with previous reports demonstrating that activity promotes the enlargement of spines^66^. This was accompanied by a concomitant reduction in the fraction of intermediate spines, while the fraction of filopodia was unchanged (**Figure 6O-R**). In contrast, the proportion of mushroom spines was similar between EE and SH conditions in both *Gfap Smo* CKO and *Gli1 Smo* CKO mice (**Figure 6N, Supplemental Figure 5**) indicating that selective perturbation of Shh signaling in astrocytes interferes with activity-dependent spine enlargement. These mice also showed no difference in the fraction of spines with intermediate or filopodial morphology between EE and SH conditions (**Figure 6P, R, Supplemental Figure 5**). These data suggest that activity-dependent Hevin promotes the enlargement of spines in a Shh-dependent manner.

## DISCUSSION

In this study, we demonstrate that neural activity stimulates Shh signaling between neurons and astrocytes. Activity of the pathway is highly dynamic and responsive to both an increase and decrease in neural activity, key features for effective translation of experience into functional gene expression. Neural activity also promotes expression of the astrocyte-derived synapse modifying molecules, Hevin and SPARC, in a Shh-dependent manner. Selective perturbation of the pathway in astrocytes occludes experience-dependent structural plasticity in deep, but not upper, layer cortical synapses, demonstrating highly localized regulation over astrocyte modulation of synapses. Taken together, these data demonstrate that neural activity stimulates Shh signaling and identify the Shh pathway as a molecular mechanism by which astrocytes convert neuronal activity into gene expression, promoting synaptic plasticity.

Transcriptional profiling demonstrates that activity induces gene expression changes in astrocytes that include early and late response genes as well as genes associated with synapse modification^2,4^. Activity stimulates dynamic Ca^2+^ signals^67^ and intracellular cAMP signaling promotes synaptic plasticity^68^. Yet the precise molecular mechanisms by which astrocytes translate neural activity into changes in gene expression have not been identified. Here we show that neural activity bidirectionally modulates Shh signaling between neurons and astrocytes in the cortex. Neural activity generated by either chemogenetic or environmental stimulation increased Shh signaling and conversely, sensory deprivation or removal of the stimulus downregulated the pathway. Notably, activity-dependent Shh signaling was selectively increased in appropriate circuits as somatosensory stimulation failed to increase Shh signaling in the visual cortex. While EE increases the number of astrocytes transducing SHH, neurons expressing *Shh* remains consistent between EE and SH conditions. This suggests that Shh signaling is initiated by a defined population of neurons that can modulate the number of astrocytes they communicate with depending on need. In this way, Shh signaling cooperates with glutamatergic signaling, acting as a molecular mechanism by which neurons recruit astrocytes during times of heightened activity promoting astrocyte interactions with synapses.

Shh signaling is a pleiotropic molecular signaling pathway with diverse roles in CNS development and function. Though best characterized for its roles as a morphogen and mitogen during embryonic neural development, several lines of evidence point to a role for SHH in regulating synapses. Genetic activation of the pathway selectively in astrocytes increases cortical synapse density that is mediated through upregulation of synapse-modifying genes^34^. Consistent with this, our data show that the synaptogenic cues, Hevin and SPARC, are regulated by Shh signaling. Interestingly, selective loss of Shh signaling in astrocytes from birth also increases synapse number. However this phenotype arises due to perturbation of developmental elimination of cortical synapses, leading to elevated synapse number that persists into adulthood^35^. The experiments in this study were performed on mice at P21, corresponding to the end of critical period plasticity for the somatosensory cortex^69^, suggesting that Shh signaling also mediates astrocyte modulation of mature, relatively established circuits. Collectively, these observations demonstrate that SHH regulates gene expression programs that promote astrocyte modulation of synapses in both developing and mature circuits. In addition, our observation that Shh signaling can be regulated by neural activity uncovers a novel dimension to this versatile signaling pathway that exerts powerful control over diverse cellular behaviors from development into adulthood.

Astrocyte modification of synapses is mediated by a diverse repertoire of molecular cues, including the release and secretion of matricellular and cell adhesion molecules^70,71^. Constitutive activation of Shh signaling in cortical astrocytes increases SPARC expression^34^, suggesting SHH is sufficient for its upregulation. Here, we extend this finding and report that selective interruption of pathway activity in astrocytes significantly decreases SPARC expression, demonstrating a requirement for Shh signaling in regulating SPARC. It has been demonstrated that SPARC promotes GluA1 AMPARs at the synapse. Although we failed to observe an increase in GluA1 in spines following EE, despite an increase in SPARC, it is possible that the levels of SPARC induced by activity in this paradigm are insufficient to promote GluA1 AMPARs at the synapse. Indeed, only intermediate concentrations of SPARC were sufficient to increase synaptic GluA1 AMPARs in culture^63^. Further experiments are necessary to determine whether a stronger stimulus or more prolonged exposure to sensory enrichment promotes a greater increase in SPARC expression and concomitant GluA1 AMPARs.

Nevertheless, we found that exposure to EE increases Hevin, a structurally related protein to SPARC with synaptogenic function, in a SHH-dependent manner. Immunohistochemical labeling, together with genetic marking of *Gli1*-expressing astrocytes, shows that both Hevin and SPARC are enriched in cells with Shh activity. However, it should be noted that upper layer astrocytes also express these proteins, suggesting that their expression is regulated by additional mechanisms that remain to be discovered. Hevin promotes the structural formation of synapses through its interactions with neurexin and neuroligin and also promotes spine maturation^61,72^. Our data show that both spine density and the proportion of spines with a mushroom morphology increase in a SHH-dependent manner following sensory experience. Taken together, these observations are consistent with the idea that experience-dependent structural plasticity is mediated by upregulation of Hevin in a SHH-dependent manner. In contrast to Hevin, the actions of SPARC on synapses are less well understood. SPARC has been shown to both antagonize and promote synapses^26,63^. Our data showing that synapse density increases following sensory experience, despite an increase in SPARC expression, would seem to be in conflict with reports that SPARC antagonizes the synaptogenic actions of Hevin. An intriguing possibility is that Hevin and SPARC are differentially upregulated at specific synapses, enabling local regulation by specific perisynaptic astrocyte processes. In support of this, shRNA knockdown of SPARC in astrocytes selectively reduces cortico-cortical VGlut1-labeled synapses, but not thalamocortical VGlut2-labeled synapses in deep layers^34^. Both the Hevin and SPARC antibodies used in this study were raised in the same species, precluding a direct test of this possibility, however future studies including EM approaches are necessary to fully elucidate the role of SPARC in SHH-dependent astrocyte modulation of synapses.

Although active Shh signaling is found predominantly in deep cortical layers, several components of the pathway, including *Gli2*, *Gli3,* and *Ptch1* are found in upper layer astrocytes^32^, suggesting these cells fail to receive sufficient ligand to initiate pathway activity. This suggests that promoting SHH availability to upper layer astrocytes would promote similar effects on synapses. In support of this, ectopic expression of *Shh* in upper layer neurons promotes expression of SHH target genes, including *Sparc* and *Kcnj10*^34^. The precise mechanism restricting SHH to deep layers is not known. In development, the SHH gradient is established by interactions with extracellular matrix (ECM) proteins, including heparin sulfate proteoglycans (HSPGs)^73^. In *Drosophila*, the movement of Hedgehog (Hh) across cells is regulated by Dally and Dally-like, two glypicans that colocalize with Hh and act to restrict its movement^74^. Recent work in vertebrate cells suggests that loss of glypican 5 (*gpc5*) broadens the spread of SHH^73^, suggesting that glypican-mediated regulation of SHH movement is conserved. The related glypicans, glypican 4 and 6, are expressed by astrocytes and regulate AMPA receptor composition at the synapse^23^ and more recently, *gpc5* has been shown to regulate GluA2 at the synapse and promote synapse stabilization^75^. This suggests that Shh signaling and glypicans work cooperatively in astrocyte modulation of synaptic function.

Precisely how SHH is released from neurons is poorly understood. Several lines of evidence point to axonal trafficking and release^76,77^, consistent with activity-dependent release. Studies in cell culture and by EM demonstrate that SHH is localized with synaptic vesicles^44^ and is released by high frequency stimulation^39^. Although cells with *Gli1* activity are found most abundantly in layers IV and V of the cortex, we found few *Shh*-expressing neurons in the thalamus. Instead, *Shh* expression is found predominantly in layer V pyramidal neurons, suggesting that Shh activity in the cortex is derived from local neuronal sources, rather than long distance projections. This may arise from recurrent collateral innervation by layer IV and V neurons^52^. SHH is also found in extracellular vesicles^78,79^, specialized vesicles derived from endosomes carrying diverse cargoes, including proteins, lipids and RNA molecules, that mediate cell-cell communication. Exosome release is stimulated by neuronal activity^80^. The localization of *Gli1*-expressing cells in deep cortical layers, adjacent to *Shh*-expressing soma, is consistent with an exosomal source of SHH. These mechanisms are not mutually exclusive and further work with SHH fusion proteins to identify its localization within specific cellular compartments could shed light on the precise mechanism by which neurons release SHH. Interestingly, our genetic labeling experiments demonstrated that activity did not increase the number of neurons expressing *Shh*. This suggests that activity increases the number of synaptic vesicles or exosomes releasing ligand, enabling greater availability in the extracellular space to reach more cells.

Here, we focus on Shh activity in the somatosensory cortex. However, Shh signaling is found throughout multiple cortical regions, including in the auditory and visual cortex, as well as in the frontal cortex^32^. In contrast, Shh signaling in the hippocampus is largely absent in astrocytes^32,35^ and instead is found predominantly in adult neural stem cells^81^, where its activity is required for their maintenance^82,83^. Notably, differential activity of the pathway is also found within a given region. Our data show that genetic deletion of Shh signaling, selectively in layers IV and V, does not perturb structural plasticity in layer II/III, suggesting that astrocyte modulation of synapses is mediated by distinct mechanisms, even within a defined circuit. This suggests that Shh signaling is broadly available to multiple, discrete circuits as a molecular mechanism by which astrocytes convert neural activity into changes in gene expression to support synaptic function and plasticity.

## Acknowledgements

We gratefully acknowledge Dr. Corey Harwell and Dr. Lucas Cheadle for thoughtful comments on this manuscript and throughout the project. The authors would also like to acknowledge Pooja Sakthivel for technical support on this project, as well as all members of the Garcia lab. This research was supported by NIH grants R21NS116664 and R01NS096100.

## Author Contributions

Conceptualization, A.D.L., M.F., A.D.R.G.; Investigation, A.D.L., M.F., R.K., G.Z.; Writing A.D.L. and A.D.R.G.; Review and editing, A.D.L. and A.D.R.G.; Funding acquisition, A.D.R.G.; Supervision A.D.R.G.

## Key Resources Table

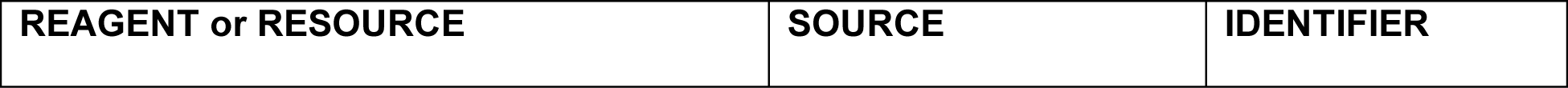
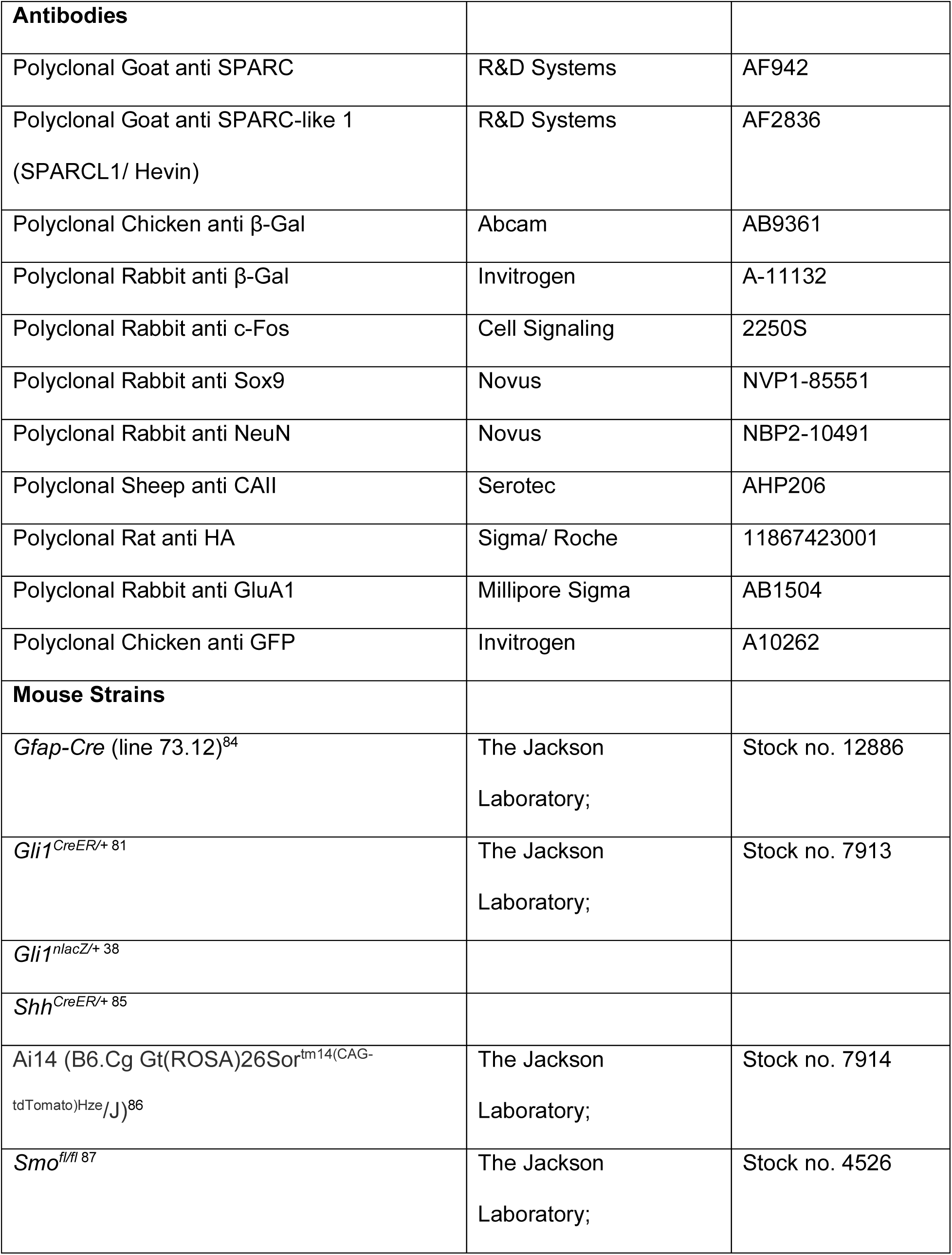
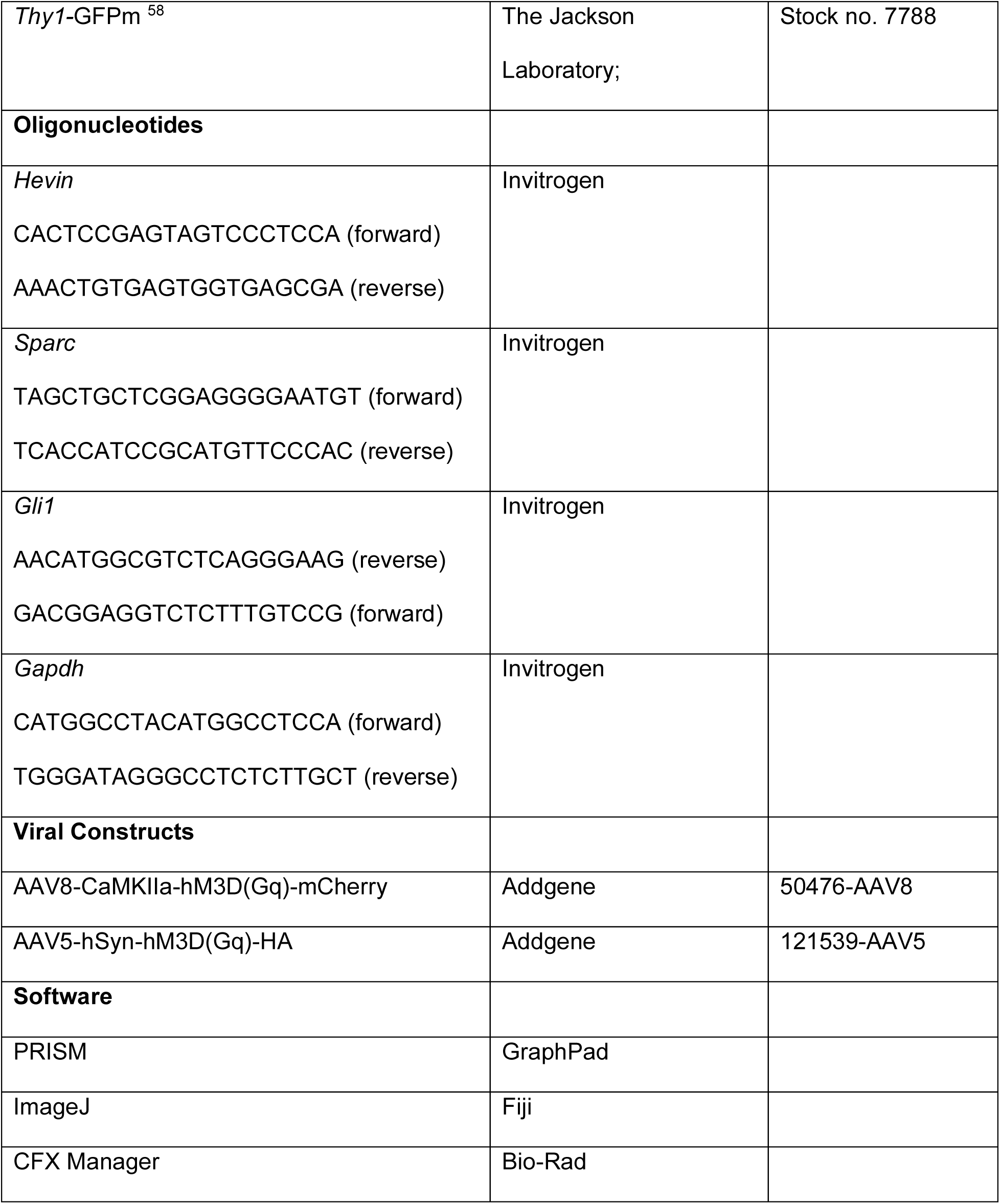

## METHODS

### Animals

All experiments were approved by Drexel University’s Institutional Animal Care and Use Committee and were conducted according to approved protocols. Transgenic mouse lines are listed in the key resources table, used on the C57BL/6 background. Postnatal (P) day 18 – 90 mice were used for this study. Mice of both sexes were included in each experiment.

### Enriched experience housing

P21 littermates were weaned into either enriched environment (EE) or standard housing (SH) cages for two days or three weeks. Large rat cages were used. The EE cage contained strings of beads hung from wire lids, positioned densely throughout the cage, requiring mice to constantly interact with the beads. The density and spacing of the beads were altered every 3 days (in 3-week experiment) to promote novelty. The SH cage contained only standard hub and nesting materials. A minimum of 7 mice was used for each cage. Data from each analysis was collected over 2-5 separate experiments from separate litters.

### Whisker trimming

Whisker trimming procedure was conducted on P21 mice. Mice were anesthetized with isoflurane in an induction chamber (3%) and then placed under dissecting scope with continuous isoflurane inhalation (1.5%). Micro scissors were used to trim whiskers on the right side, as close to the skin as possible.

Whiskers were trimmed every 3 days for the duration of 3 weeks.

### Tamoxifen administration

Tamoxifen (Sigma) was dissolved in corn oil at a 20 mg/ml concentration at 37°C. For *Gli1^CreER/nlacZ^*; Ai14 mice (Figure 2), 3 doses of 250mg/kg tamoxifen were administered by oral gavage over 3 days from P18-P20 and tissues were analyzed at P42. For *Shh^CreER^*; Ai14 mice (Figure 4), 3 similar doses were administered over 3 days at the end of the experiment (concurrent with when increases of BGal cells were observed). For *Gli1^CreER^*; *Smo^fl/fl^;* Ai14 mice (Figure 5, 6), 3 similar doses were administered from P18-P20, weaned into SH or EE cages from P21-P23 and analyzed on P23.

### Virus injection

All AAV injections were conducted on adult mice at P60 or older. Immediately prior to surgery, mice received a subcutaneous injection of carprofen (Rimadyl, 6.8 mg/kg), and a follow up injection given the day after surgery. Mice were anesthetized with isoflurane in an induction chamber (3%) and then placed in a stereotaxic apparatus (Stoelting) with continuous isoflurane inhalation (1.5%). The scalp was opened and a small area of the skull was removed near injection site. Stereotaxic coordinates for somatosensory cortex injection were AP -1.5 mm, ML -2.5 mm, DV -0.5 mm.

Viruses were stored in aliquots at -80°C and kept on ice until injection. Virus titers were ≥ 3×10¹² vg/mL for AAV8-CaMKIIa-hM3D(Gq)-mCherry and ≥ 7×10¹² vg/mL for AAV5-hSyn-hM3D(Gq)-HA. Virus was front-filled into a glass syringe (Hamilton). Injection volume was 1000 nL, delivered at 200 nL/ min controlled by microinjector (WPI). After injection, mice were allowed to recover for 2 weeks before administration of CNO (Tocris). CNO was dissolved in drinking water, aimed to deliver a dosage of 1mg CNO per kg mouse over 1 week. Control mice received water only.

### Collection of animal tissue

Animals used in experiments were each assigned a unique 4-5 digit ID in order to blind researchers of genotypes and housing conditions. Mice were deeply anesthetized by intraperitoneal injection of ketamine/ xylazine/ acepromazine, transcardially injected with 100 units heparin and perfused with ice-cold PBS followed by 4% paraformaldehyde (PFA; Sigma-Aldrich). Brains were dissected and fixed for at least 4 hours in PFA at 4°C then transferred to 30% sucrose at 4°C for at least 48 hours. Brains were cryosectioned (Leica CM3050s) and collected in 40 µm sections. Sections were stored in 0.1M Tris-buffered saline (TBS) with 0.05% sodium azide at 4°C. Immunohistochemistry was performed with primary antibodies listed in the resources table below. Sections were washed three times with 0.1M TBS at room temperature (10 minutes/wash) then blocked in 10% Normal Serum with 0.5% Triton-X (Sigma-Aldrich) for one hour at room temperature. Sections were then incubated with primary antibody in 0.5% Triton-X at 4°C overnight.

### Immunohistochemistry

For fluorescent immunohistochemistry, sections were rinsed for three times the following day in 0.1M TBS and incubated in AlexaFluor-conjugated secondary antibodies with 10% Normal Serum in 0.1M TBS at room temperature for two hours. Sections were then rinsed in 0.1M TBS and incubated in DAPI (Life Technologies) for 15 minutes. Sections were rinsed again in 0.1M TBS, mounted on microscope slides (Fisherbrand), and coverslipped using ProLong Gold Antifade Mounting Medium (Invitrogen #P10144) and Fisherfinest Premium Cover Glass. For brightfield immunohistochemistry, sections were rinsed as previously described but were incubated with biotinylated secondary antibodies against the primary species (Vector Laboratories) for one hour. Sections were transferred to Avidin-Biotin Complex solution (Vector) for one hour and visualized using 3,3’-diaminobenzadine (DAB Peroxidase Substrate Kit, Vector). Sections were then mounted onto slides as described, dehydrated through increasing concentrations of EtOH, and cover slipped with DPX mounting medium (Fisher).

### Fluorescent In situ hybridization (RNAscope)

All fluorescent in-situ hybridization (FISH) experiments were performed on brain tissue fixed with 4% paraformaldehyde and processed using cryosectioning at 14 µm and directly slide mounted, with one brain section per slide. The assay was carried out using the “fixed frozen sample preparation and treatment” and the RNAscope Multiplex Fluorescent v2 Assay using the manufacturer’s instructions. The original protocol can be found here on the ACDBio website: https://acdbio.com/documents/product-documents.

Confocal z-stack images were acquired at 63x for blinded analysis in ImageJ (FIJI). Cells with at least 4 puncta were identified as positive for *Gli1* mRNA. The number of *Gli1+* cells were divided by the total number of DAPI cells within the z-stack, and then normalized to SH condition for each layer. For each animal, each layer, at least 2 z-stacks were analyzed for a minimum of 100 DAPI cells.

### Stereological quantification

The number of cells in each area of the cortex was estimated using a modified optical fractionator and stereological image analysis software (Stereo Investigator, MBF Bioscience) with upright microscope (Zeiss) with a computer-driven stage. The cortical area of interest (somatosensory, motor or visual cortex) was outlined at a low magnification, using anatomical landmarks as described in Paxinos and Franklin’s The Mouse Brain in Stereotaxic Coordinates 4th Ed. For somatosensory cortex, sections from A-P Bregma 1.33mm to -2.15mm were analyzed. For visual cortex, sections from A-P Bregma -2.27mm to -4.59mm were analyzed. A target cell count of 150 and target coefficient of error (m=1) of < 0.1 was used to define number of sections counted, scan grid, and counting frame size. Counting frames were randomly selected by the image analysis software and cells were counted at the 40x objective with a DIC optic filter. Only cells with a clearly labeled cell body were counted.

### Fluorescent intensity analysis

Immunofluorescent slides were imaged using Leica DMI 4000B microscope equipped with TCS SPE confocal system. Regions of interest were identified with 10x dry objective and confocal z-stacks were acquired with 40x oil objectives. To analyze protein abundance in a single astrocyte, a 20 μm x 20 μm region of interest is drawn in ImageJ around an astrocyte identified by fluorescent signal of either tdTomato reporter protein or staining for βGal or Sox9. This region of interest should contain one astrocyte of interest and not contain signal from other cells. The mean fluorescent intensity is then collected. At least 25 cells were sampled per animal, dispersed throughout multiple sites encompassing the somatosensory cortex. The mean fluorescent intensities of the cells were averaged to produce the fluorescent intensity value for one animal, represented in the graph as one data point. For negative controls, we measured fluorescent signals of Sox9 and found no difference between layers and housing conditions.

### Dendritic spine analysis

Analysis of spine density was performed in a blinded study design. WT controls were derived from Cre-negative littermates. For analysis in the cortex, apical dendrites of layer V pyramidal neurons located in layers II/III, IV and V were imaged. For analysis in the hippocampus, apical dendrites from CA1 pyramidal neurons were imaged. Confocal z-stacks were collected with 63x oil objective, at 0.35 μm steps. Spine density was determined by first tracing the dendritic segment in ImageJ and obtaining the length. Analyzed segments range from 75 – 125 µm in length. Protrusions were classified into mushroom, intermediate or filopodial morphologies. Protrusions exhibiting a bright, bulbous head with a large head-to-stalk ratio were classified as mushroom. Thinner protrusions with a dimmer, smaller or no head were classified as intermediate. Dim, thin and elongated protrusions that lack any head were classified as filopodia. For GluA1 colocalization analysis in dendritic spines, 63x z-stack images were analyzed. Only spine heads colocalizing with a clearly defined GluA1 puncta were categorized as positive for GluA1. The number of GluA1 positive protrusions were divided by the total number of protrusions on the segment to obtain the fraction of GluA1 positive protrusions.

### Quantitative reverse transcriptase – polymerase chain reaction (qRT-PCR)

Mice were deeply anesthetized by isoflurane before rapid decapitation. Cortices were dissected into TRIzol reagent (Thermo-Fisher, Waltham, MA) and RNA was extracted according to TRIzol’s standard protocol. RNA was reverse transcribed to cDNA using the High-Capacity cDNA Reverse Transcription Kit (Thermo-Fisher). qPCR was performed with the CFX96 Touch Real-Time PCR Detection System (Bio-Rad) using PowerUp SYBR Green Master Mix (Thermo-Fisher). Primers were designed using NCBI Primer-BLAST (see resources table). Samples were run in triplicate. Data was analyzed using the CFX Manager Software (Bio-Rad), using the ΔΔCt method. *Gapdh* was used as the reference gene, itself having no difference of expression between sample groups.

### Quantification and statistical analysis

Statistical analysis and data visualization were performed using Prism (GraphPad). Data were assessed for normality using Shapiro-Wilk test. For t-tests, variances were compared before choosing between Student’s (if no significant difference) or Welch’s t-tests (if there is a significant difference).

Pair-wise t-tests were used for measurements taken from the same animal. Additionally, one-way and two-way ANOVA with Tukey’s multiple comparisons were used with reported p-values. In all graphs, data points represent individual animals.

## Supplementary

**Figure S1: Enriched experience stimulates Shh activity in astrocytes. Related to Figure 2**.

(**A-D**) Immunofluorescence for βGal (green), and Sox9 (A), NeuN (B), CAII (C) or Ki67 (D; magenta) in the cortex of *Gli1^nlacZ/+^* mice. Counterstained with DAPI (blue). Merged images shown in right panels. (**E**) The fraction of βGal-labeled cells colocalizing with Sox9 does not change between SH and EE. *n=4* mice in SH, *n=4* mice in EE, 100-400 βGal cells analyzed per animal. Scale bars, 10 μm; data points represent individual animals; bars show mean ± SEM; Student’s t-test.

**Figure S2: Whisker deprivation reduces SHH activity.**

(**A-B**) Sterelogical quantification of βGal cells from the ipsilateral (intact) and contralateral (deprived) barrel (A) and auditory (B) cortex after 3 weeks of unilateral whisker trimming. *n=3* mice, data points represent individual animals, bars show mean ± SEM; paired t-tests.

**Figure S3: Shh activity is effectively disrupted in *Gli1 Smo* CKO mice without altering spine density. Related to Figure 5**.

(**A-B**) qPCR for *Gli1* expression in WT and *Gfap Smo* CKO (A) and *Gli1 Smo* CKO (B) mice. *n=4-6* mice. (**C**) Representative dendritic segments from P60 WT versus *Gli1 Smo* CKO at standard housing conditions. Scale bar, 5 μm. (**D**) Protrusion density of deep layer dendrites between P60 WT vs *Gli1 Smo* CKO at standard housing conditions. *n=2* WT mice, *n=3 Gli1 Smo* CKO mice, 3 dendritic segments analyzed per animal. (**E**) Spine density of deep layer dendritic segments in wild-type littermate controls of *Gli1 Smo* CKO animals housed in SH, including Cre-, Cre- / + tamoxifen, Cre+ (no tamoxifen). Animals were subsequently pooled as WT for comparison with *Gli1 Smo* CKO. (**F**) Spine density of deep layer dendritic segments in wild-type littermate controls of *Gli1 Smo* CKO animals housed in EE, including Cre- and Cre+ (no tamoxifen). Animals were subsequently pooled as WT for comparison with *Gli1 Smo* CKO. Data points represent individual animals; bars show mean ± SEM; statistical analyses were Welch’s t-test (A), Student’s t-tests (D, F) and one-way ANOVA with Tukey’s multiple comparisons (B, E).

**Figure S4: Hevin and SPARC are regulated by Shh activity. Related to Figure 6**.

(**A, C**) Immunostaining for Hevin (gray, A) and SPARC (gray, C) in the cortex of P60 *Gli1^CreER/+^;* Ai14 mice showing tdTom cells (magenta) primarily localized to deep layers (IV and V) of the cortex in contrast to upper layer (II/III). (**B, D**) Fluorescent intensity measurements of Hevin (B) and SPARC (D) across cortical layers. *n=5-8* animals, 2 sections analyzed per animal. (**E, G**) Immunofluorescent staining for Hevin (E, gray, left panels) and SPARC (G, gray, left panels) in Sox9 cells (green) from *Gli1 Smo* CKO mice housed in SH vs EE. Merged images in right panels. Scale bar, 10 μm. (**F, H**) Fluorescent intensity analysis of Hevin (F) and SPARC (H) in Sox9 astrocytes from *Gfap Smo* CKO mice housed in SH or EE from P21-P23. Data points represent individual animals; bars show mean ± SEM; statistical tests are one-way ANOVA with Tukey’s multiple comparisons (B, D) and Student’s t-tests (H, J).

**Figure S5: Enriched experience does not increase GluA1 AMPAR in dendritic spines. Related to Figure 5 and 6**.

(**A - C**) Fraction of mature (A), intermediate spines (B) and filopodia (C) in *Gfap Smo* CKO mice housed in SH or EE. (**D**) Immunostaining for GluA1 (magenta) in dendritic spines (gray). Arrow points to spine colocalizing with GluA1 puncta. Merged images in right panel. Scale bar, 5 μm. (**E**) The fraction of GluA1+ protrusions in WT, *Gfap Smo* CKO and *Gli1 Smo* CKO mice after SH or EE. At least 200 spines analyzed per animal. Data points represent individual animals; bars show mean ± SEM; statistical tests are Student’s t-tests (A, B, C) and two-way ANOVA with Tukey’s multiple comparisons (E).

