## Supplemental Figures for "Astrocyte modulation of synaptic plasticity mediated by activity-dependent Sonic hedgehog signaling"

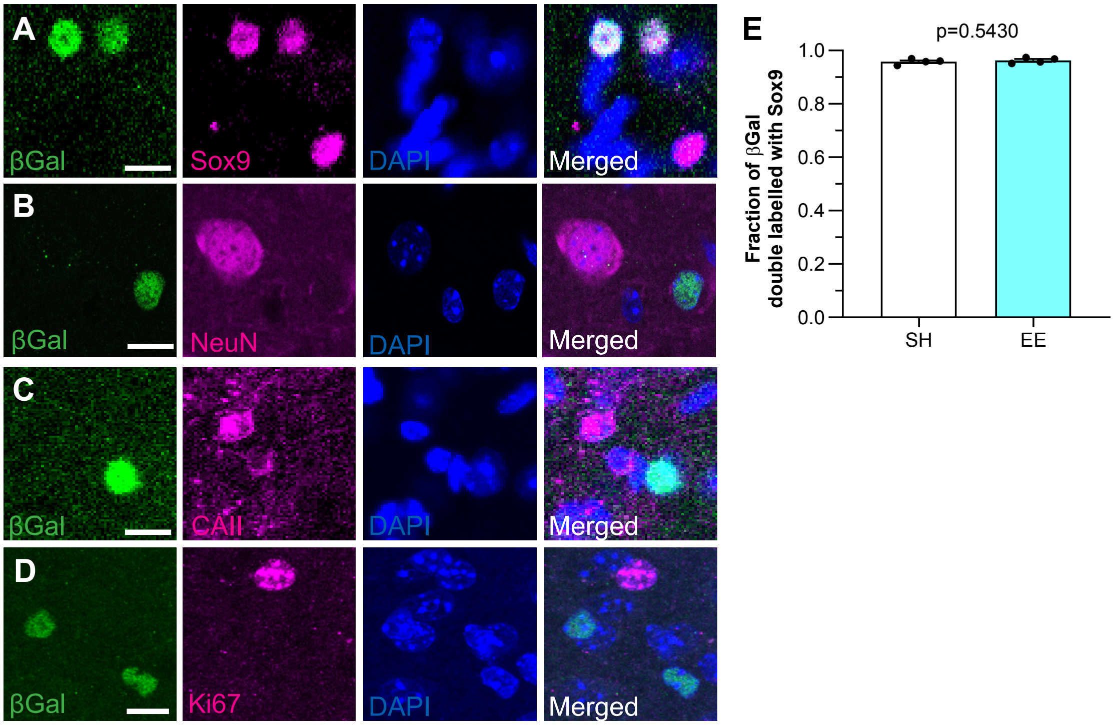


**Figure S1: Enriched experience stimulates Shh activity in astrocytes. Related to Figure 2.**

**(A-D)** Immunofluorescence for βGal (green), and Sox9 (A), NeuN (B), CAII (C) or Ki67 (D; magenta) in the cortex of *Gli1*$^{nlacZ/+}$ mice. Counterstained with DAPI (blue). Merged images shown in right panels. **(E)** The fraction of βGal-labeled cells colocalizing with Sox9 does not change between SH and EE. *n=4* mice in SH, *n=4* mice in EE, 100-400 βGal cells analyzed per animal. Scale bars, 10 μm; data points represent individual animals; bars show mean ± SEM; Student's t-test.

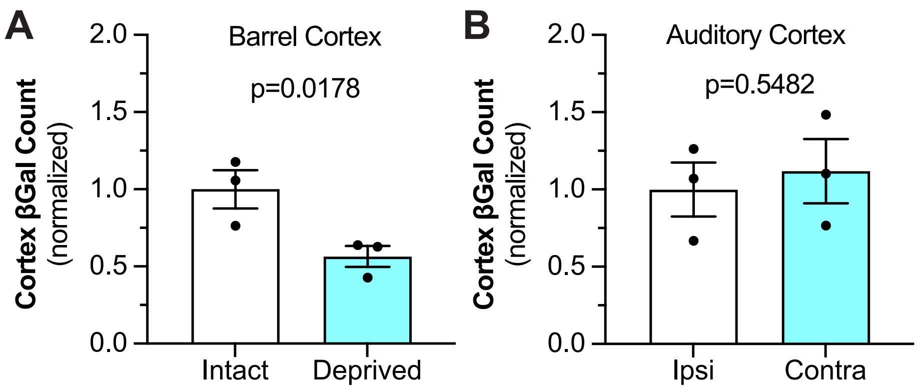


**Figure S2: Whisker deprivation reduces Shh activity.**
**(A-B)** Sterelogical quantification of βGal cells from the ipsilateral (intact) and contralateral (deprived) barrel (A) and auditory (B) cortex after 3 weeks of unilateral whisker trimming. *n=3* mice, data points represent individual animals, bars show mean ± SEM; paired t-tests.

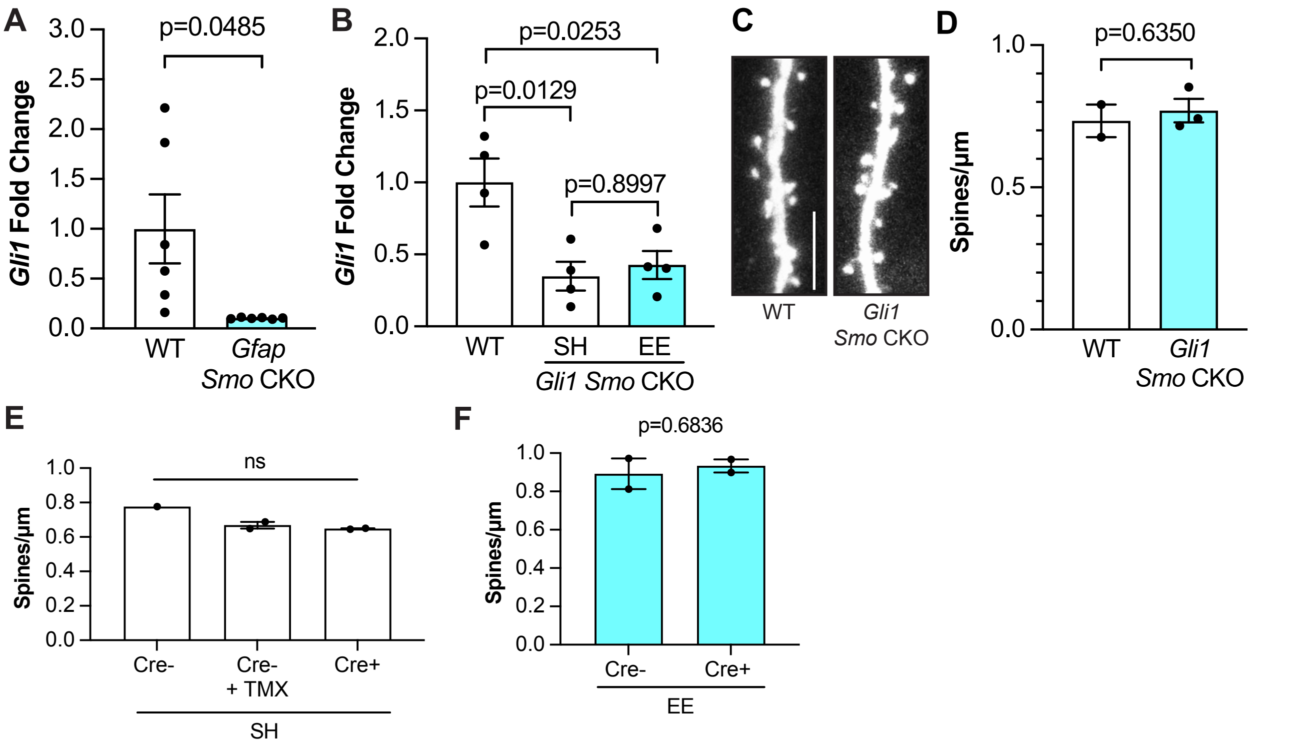


**Figure S3: Shh activity is effectively disrupted in *Gli1 Smo* CKO mice without altering spine density. Related to Figure 5.**

**(A-B)** qPCR for *Gli1* expression in WT and *Gfap Smo* CKO (A) and *Gli1 Smo* CKO (B) mice. *n=4-6* mice. **(C)** Representative dendritic segments from P60 WT versus *Gli1 Smo* CKO at standard housing conditions. Scale bar, 5 µm. **(D)** Protrusion density of deep layer dendrites between P60 WT vs *Gli1 Smo* CKO at standard housing conditions. *n=2* WT mice, *n=3 Gli1 Smo* CKO mice, 3 dendritic segments analyzed per animal. **(E)** Spine density of deep layer dendritic segments in wild-type littermate controls of *Gli1 Smo* CKO animals housed in SH, including Cre-, Cre- / + tamoxifen, Cre+ (no tamoxifen). Animals were subsequently pooled as WT for comparison with *Gli1 Smo* CKO. **(F)** Spine density of deep layer dendritic segments in wild-type littermate controls of *Gli1 Smo* CKO animals housed in EE, including Cre- and Cre+ (no tamoxifen). Animals were subsequently pooled as WT for comparison with *Gli1 Smo* CKO. Data points represent individual animals; bars show mean ± SEM; statistical analyses were Welch's t-test (A), Student's t-tests (D, F) and one-way ANOVA with Tukey's multiple comparisons (B, E).

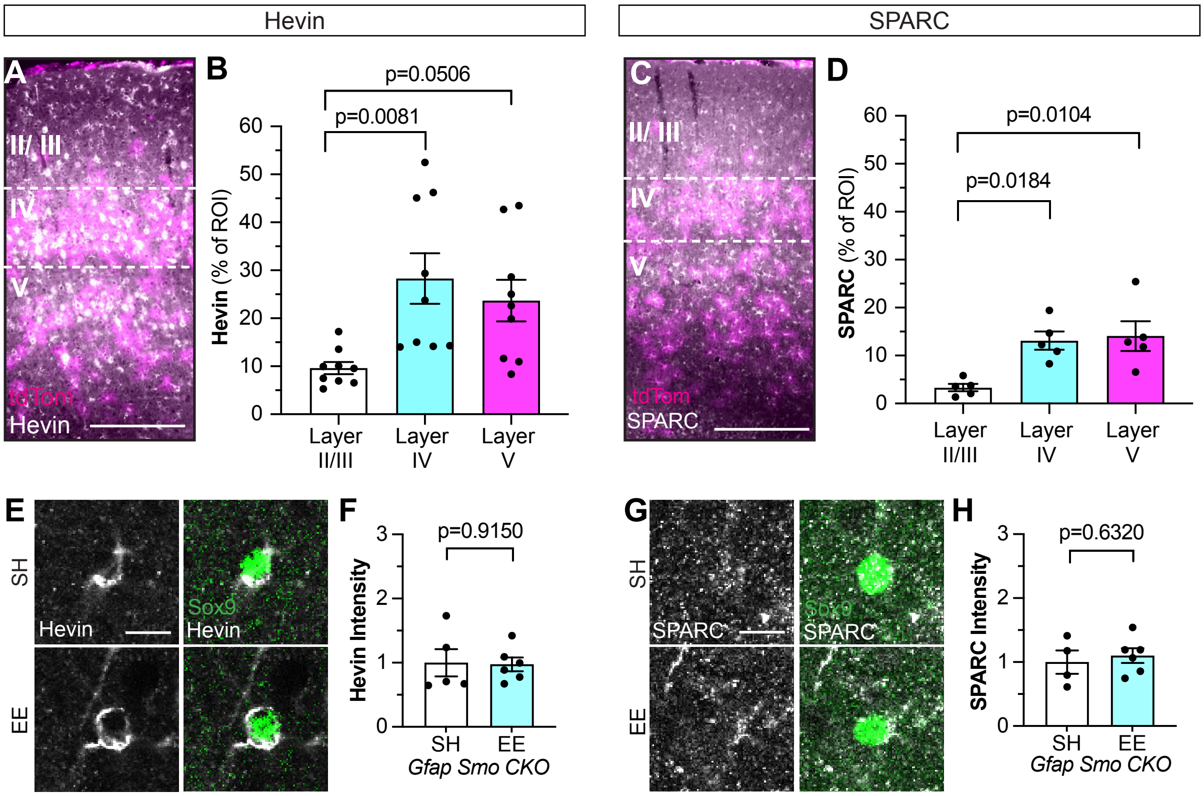


**Figure S4: Hevin and SPARC are regulated by Shh activity. Related to Figure 6.**
**(A, C)** Immunostaining for Hevin (gray, A) and SPARC (gray, C) in the cortex of P60 *Gli1*$^{CreER/+}$*;* Ai14 mice showing tdTom cells (magenta) primarily localized to deep layers (IV and V) of the cortex in contrast to upper layer (II/III). **(B, D)** Fluorescent intensity measurements of Hevin (B) and SPARC (D) across cortical layers. *n=5-8* animals, 2 sections analyzed per animal. **(E, G)** Immunofluorescent staining for Hevin (E, gray, left panels) and SPARC (G, gray, left panels) in Sox9 cells (green) from *Gli1 Smo* CKO mice housed in SH vs EE. Merged images in right panels. Scale bar, 10 μm. **(F, H)** Fluorescent intensity analysis of Hevin (F) and SPARC (H) in Sox9 astrocytes from *Gfap Smo* CKO mice housed in SH or EE from P21-P23. Data points represent individual animals; bars show mean ± SEM; statistical tests are one-way ANOVA with Tukey's multiple comparisons (B, D) and Student's t-tests (H, J).

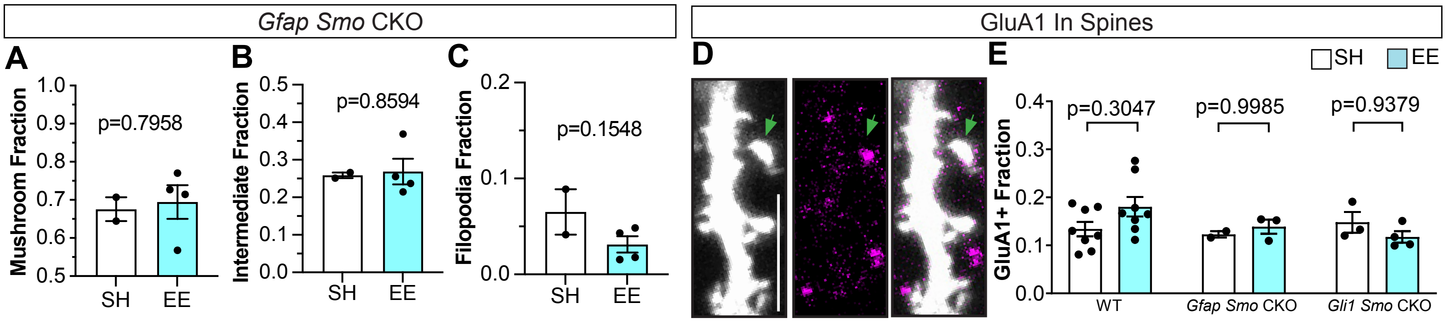


**Figure S5: Enriched experience does not increase GluA1 AMPAR in dendritic spines. Related to Figure 5 and 6.**

**(A - C)** Fraction of mature (A), intermediate spines (B) and filopodia (C) in *Gfap Smo* CKO mice housed in SH or EE. **(D)** Immunostaining for GluA1 (magenta) in dendritic spines (gray). Arrow points to spine colocalizing with GluA1 puncta. Merged images in right panel. Scale bar, 5 μm. **(E)** The fraction of GluA1+ protrusions in WT, *Gfap Smo* CKO and *Gli1 Smo* CKO mice after SH or EE. At least 200 spines analyzed per animal. Data points represent individual animals; bars show mean ± SEM; statistical tests are Student's t-tests (A, B, C) and two-way ANOVA with Tukey's multiple comparisons (E).
